# Structured hierarchical models for probabilistic inference from perturbation screening data

**DOI:** 10.1101/848234

**Authors:** Simon Dirmeier, Niko Beerenwinkel

## Abstract

Genetic perturbation screening is an experimental method in biology to study cause and effect relationships between different biological entities. However, knocking out or knocking down genes is a highly error-prone process that complicates estimation of the effect sizes of the interventions. Here, we introduce a family of generative models, called the *structured hierarchical model* (SHM), for probabilistic inference of causal effects from perturbation screens. SHMs utilize classical hierarchical models to represent heterogeneous data and combine them with categorical Markov random fields to encode biological prior information over functionally related biological entities. The random field induces a clustering of functionally related genes which informs inference of parameters in the hierarchical model. The SHM is designed for extremely noisy data sets for which the true data generating process is difficult to model due to lack of domain knowledge or high stochasticity of the interventions. We apply the SHM to a pan-cancer genetic perturbation screen in order to identify genes that restrict the growth of an entire group of cancer cell lines and show that incorporating prior knowledge in the form of a graph improves inference of parameters.

## 1. Introduction

Probabilistic graphical models (PGMs) and Bayesian hierarchical models (HMs) are integral parts of statistical data analysis and computational biology. PGMs graphically represent the joint probability distribution of several random variables by encoding conditional dependencies between variables as edges in a graph (Koller and Friedman, 2009; Maathuis et al., 2018). Bayesian HMs are a special case of PGMs, where the graph is usually a directed tree and parameters of interest are endowed with distributions which themselves are parametrized leading to a conditionally independent hierarchy of variables (Gelman et al., 2013).

In bioinformatics and computational biology, PGMs have been found to be especially useful and have a long history of applications, because biological measurements, such as gene expression values or metabolite concentrations, are often correlated making them suitable for joint probabilistic modelling. Simple PGMs, such as hidden Markov models (Durbin et al., 1998; Baldi et al., 1994; Eddy, 1998; Stanke and Waack, 2003; Marioni, Thorne and Tavaré, 2006; Yoon, 2009; Finn, Clements and Eddy, 2011) have been discussed in the bioinformatics literature at length.

Bayesian networks (BNs), a more general class of graphical models, where the underlying graph is a directed acyclic graph (DAG), are computationally more expensive, but probabilistically more expressive. For instance, Kuipers et al. (2018) used BNs to identifiy gene-gene interactions in cancer tissues and for clustering the mutational profiles of cancer types. Other approaches include Jansen et al. (2003) who used BNs to detect protein-protein interactions (PPIs) from genomic data, such as gene expression values, i.e., the abundance of messenger RNA (mRNA) of a gene in the cell, Friedman et al. (2000) where BNs were applied to model gene interactions and analyze expression data, or Sachs et al. (2005) who used BNs to reconstruct signalling networks. For time-series data, dynamic Bayesian networks (DBNs) have been used to, for instance, to identify gene regulatory networks (Murphy et al., 1999; Zou and Conzen, 2004; Li et al., 2011; de Luis Balaguer and Sozzani, 2017).

In contrast to BNs, Markov random fields (MRFs) use undirected edges to encode conditional dependencies. Wei and Li (2007) and Chen, Cho and Zhao (2011) used MRFs to encode biological pathways and successfully identified single genes related to diseases. **?** used Gaussian MRFs to model the partial correlations among metabolites and showed that strong correlations often correspond to pathway interactions. Furthermore, in a recent study Schubert et al. (2019) used binary MRFs (Ising models) to model genetic interactions in order to predict epistatic loci that affect antibiotic resistance. For analysis of time-series gene expression data, Wei and Li (2008) used a spatio-temporal MRF that models the time-course of the differential gene expression stages. Recent approaches involving graphical models and network inference in biological applications have, for instance, been reviewed in Hawe, Theis and Heinig (2019).

HMs have gained wide-spread attention in modelling of biological data, on the one hand, due to their ability to represent and model the structure of many data sets, for instance, the nested structure of data, when, for multiple genes, measurements have been made in multiple conditions (e.g., tissues, cancer types, viruses, patients), and on the other hand, because biology itself is inherently hierarchical, for instance, between genes, transcripts, and proteins. Approaches utilizing hierarchical models are, for instance, presented in Fusi et al. (2014); Rakitsch et al. (2012); Zhou and Stephens (2012); Korte et al. (2012); Loh et al. (2015).

HMs can share statistical strength for parameter estimation across hierarchy levels. If we can infer parameters well in one group of the hierarchy, it can help to infer parameters well in another. In addition, for Bayesian HMs the choice of appropriate priors can induce an auto-regularizing effect, which makes HMs suitable for low sample size scenarios. A drawback of HMs is that one cannot assess statistical significance of the random effect estimates themselves, even in the frequentist setting

HMs have been found especially useful for the analysis of biological interventional data, such as genetic perturbation screens. In these screens, the variables of interest are artificially intervened on by an experimental perturbation. The intervention can either be conducted on a genomic level through loss-of-function mutations (knock-out), or the transcriptomic level by post-transcriptional gene silencing (knock-down), or post-translationally on the proteomic level. On the genomic level, the CRISPR-Cas9 system (Jinek et al., 2012; Doudna and Charpentier, 2014), where small guide RNAs (gRNAs) are used to direct a Cas9 protein to a target gene and a missense mutation is induced, has become the method of choice. One of the most frequent applications of genetic perturbation screening is measurement of downstream causal effects after the intervention. These effects can, for instance, be changes in transcript expression of certain genes when the perturbed gene was a transcription factor, or alteration of the viability of a pathogen (Rämö et al., 2014) or of a cancer cell line (Cowley et al., 2014; Aguirre et al., 2016; Meyers et al., 2017; Patel et al., 2017). Viability is usually defined and measured as the log-fold change of the number of surviving cells pre-and post-intervention. The main interest in viability screens is to find genes that upon loss-of-function induce restricted or enhanced proliferation of cells or pathogens. Therapeutically, genes that enhance the growth of cancer cell lines or pathogens are of particular interest, since these genes can serve as potential drug targets for disease treatment. Consequently, identification of genes that enhance growth for an entire group of cancer cell lines or pathogens are of even greater interest, because of their potential to serve as targets for drugs with broad-spectrum activity.

However, genetic perturbation screens are often accompanied by several persisting problems. While improvements in the design and analysis of experiments, such as the composition of nucleotides of gRNA sequences (Doench et al., 2014; Xu et al., 2015) or using appropriate noise models (Imkeller et al., 2019), have been made, genetic perturbation screens still suffer from elevated false negative and false positive rates and bias of estimated effect sizes of the knock-outs (Munoz et al., 2016; Ong et al., 2017; Zhu et al., 2019; Meyers et al., 2017). This is due to the fact that, while the biochemical process of an intervention is well understood, other factors, such as off-target effects, gRNA cytotoxity, and binding affinity, can severely impact the success of an intervention (Wu et al., 2014; Doench et al., 2016). For instance, off-target effects might lead to perturbation of a gene which is non-essential, rather than the targeted gene which induces cell death. Similarly, high cytotoxicity of a gRNA might induce cell death, while an intervention in the target gene itself is neutral. Other sources of error include stochasticity of transport of a gRNA into a cell, copy number alterations in cancer, multiplicity of infection, and low sequencing depth. Hence, not only is our understanding of complex biological systems and the interactions of their entities still limited, but also our understanding of the process responsible for generating the data. Due to imperfect interventions, readouts of perturbations screens are often noisy and confounded and a good model considering all relevant covariables is challenging to determine.

In addition, biological screening historically faces another problem, namely low sample sizes due to high costs of genome-wide experimentation. Public genome-wide perturbation data, such as provided in the DepMap portal (Tsherniak et al., 2017), often only have as little as four replicates or less per intervention. Assessing causal effects and estimation of parameters is therefore difficult. In these settings, HMs are useful due to auto-regularization and borrowing of statistical strength, even though other approaches such as regularized linear models (Schmich et al., 2015) or empirical Bayesian procedures exist (Love, Huber and Anders, 2014; Robinson, McCarthy and Smyth, 2010; Li et al., 2014).

A promising extension of graphical and hierarchical modelling would be to combine the two approaches. PGMs are useful for probabilistically modelling biological prior information, e.g., in the form of networks, which can then be used for the analysis of data sets using HMs. While the idea of incorporating networks for biological data analyses is not new, (Dirmeier et al., 2017; Li and Li, 2008; Chen et al., 2012; Kim et al., 2012; Zitnik, Agrawal and Leskovec, 2018; Zamora-Resendiz and Crivelli, 2019), combining PGMs with HMs to aid and inform inference of posterior distributions has not been studied so far.

Here, we propose *structured hierarchical models* (SHMs), a family of models that allows incorporating biological prior knowledge in the form of graphs directly into probabilistic analyses. The main idea is to use a categorical MRF as latent labelling, i.e., clustering of genes, in order to probabilistically encode the pairwise relationships of genes using biological networks (Figure 1). Genes that are functionally related and are neighbors in the graph have a higher probability to belong to the same cluster than to a different one. For example, for cell-viability assays, the clustering would label genes as essential or non-essential. If a gene is labelled as non-essential then the probability of its neighbors to also be non-essential is increased and vice versa. SHMs use the clustering of genes to inform the inference of parameters in an HM for which the data generating process is extremely noisy, difficult to model, and often yields erroneous inferences of parameters due to misspecification. The MRF pushes information of gene relationships downwards to inform the inference of parameters. In comparison to other methods that cluster variables in the data space, our method clusters data in the latent space of the gene effects. By encoding interactions of genes through a MRF we incorporate structural information into the inference. Since the model is fully Bayesian, appropriate choice of priors can furthermore have an auto-regularizing effect making it especially useful for low sample size scenarios.

**Fig 1:**
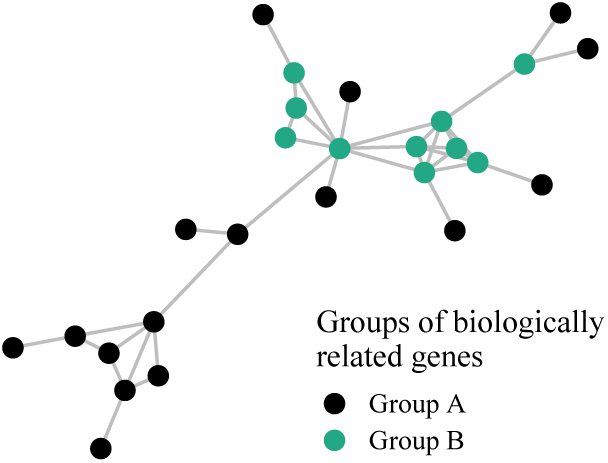
A biological network of gene-gene interactions that consists of two groups of genes, *A* and *B*. Edges define functional relationships between two genes. We use a Markov random field to encode similarities of pairs of genes, where neighboring genes have a higher probability to be in the same group than in a different one.

The rest of the paper is organized as follows: we first introduce the SHM and its basic structure, i.e., the core graphical model. We then show some empirical properties of the SHM in a study using simulated data, before we apply it to a biological data set from the DepMap portal (Tsherniak et al., 2017). The DepMap data consists of multiple different cancer cell lines for which genome-wide perturbation screens have been conducted in quadruplicate. It is a low sample size data set of multiple conditions that due to the nature of perturbations should exert high variance. Inference of essential genes in multiple conditions is of great medical interest, because it could allow divising drugs with broad-spectrum activity. We conclude the paper with some remarks about Bayesian modelling and the SHM in general.

## 2. Structured hierarchical models

Genetic perturbation screens often exhibit high noise levels and are difficult to model due to frequent incomplete domain knowledge, high stochasticity of interventions, and low sample sizes. If perturbations are conducted for multiple conditions, such as cancer cell lines, the data have a nested structure which suggests a hierarchical modelling approach. We combine the hierarchical model with a categorical Markov random field to incorporate biological prior information in the form of networks to the model. We first describe the two model components separately and then introduce the SHM.

### Bayesian hierarchical model

SHMs use Bayesian HMs as the first component to model a data set *y*_*gcn*_ with genes *g* ∈ {1, …, *G*}, conditions *c* ∈ {1, …, *C*}, e.g., cell types, patients or tissues, and observations *n* ∈ {1, …, *n*_*gc*_}. The hierarchical model for the data set *y*_*gcn*_ is defined follows:

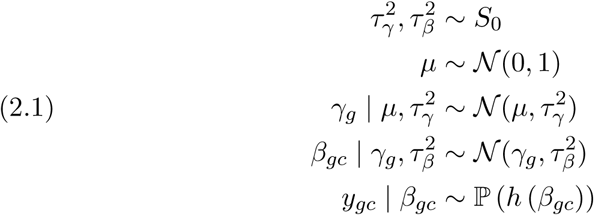

The top level of the hierarchy consists of gene effects *γ*_*g*_ which describe the impact of a perturbation of a gene on the cell. The gene effects *γ*_*g*_ are modelled as normal random variable, with mean *µ* and standard deviation *τ*_*γ*_ . The level below consists of gene effects per condition *β*_*gc*_ which are parameterized by the gene effects *γ*_*g*_ and a standard deviation *τ*_*β*_, and describe the effect of a perturbation on gene within a specific condition. We choose a unspecific joint prior distribution *S*_0_ over 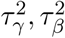 that can be specified appropriately, e.g., subjectively or by formal rules (Kas and Wasserman, 1996; Gelman, 2006; Kass et al., 2006). In the model definition above, we assume homoscedasticity of the errors 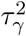 and 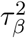, but this is not a requirement and the distributions can be adapted, too. On the lowest level the random variables *β*_*gc*_ are used to model the distribution of the data which we assume to follow and exponential family distribution ℙ. For instance, for continuous data *y*_*gcn*_ might be normally distributed, while it could follow a Bernoulli distribution in the binary case. Like in a generalized linear regression model (GLM), we use a link-function, *h*^−1^, to relate the latent variable *β*_*gc*_ to the mean of the data.

### Markov random field

The second component of an SHM is a categorical MRF **z** ∈ {1, …, *K*}^*G*^ that encodes the assignment of a gene *g* to one of *K* components, for example, clusters of essential and non-essential genes. The categorical variables **z** are distributed

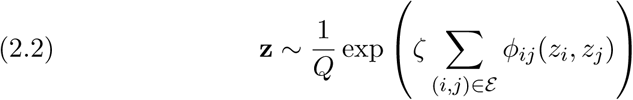

where *Q* is a normalization constant, *ζ* is a weight, and ***ε*** are the edges of a biological network, such as a PPI. The functions *ϕ*_*ij*_ are potentials defined as

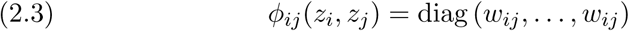

The hyperparameters *w*_*ij*_ are non-negative edge weights of the network. Thus we encode biological prior knowledge of related genes as probability distributions where two genes *i* and *j*, if biologically related, have an increased probability of having the same label *z*_*i*_ and *z*_*j*_.

### Structured hierachical model

SHMs combine Bayesian HMs and categorical MRFs, by replacing the distribution of *γ*_*g*_ in Equation (2.1) with a distribution conditioned on the MRF. The entire model is defined as follows (Figure 2):

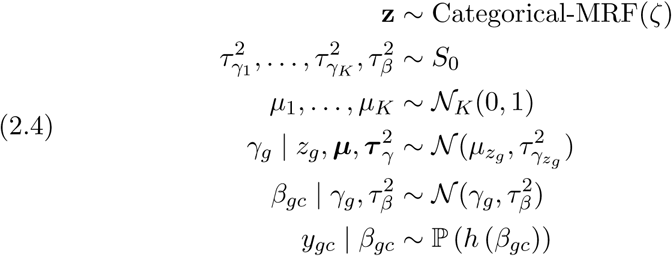

**Fig 2:**
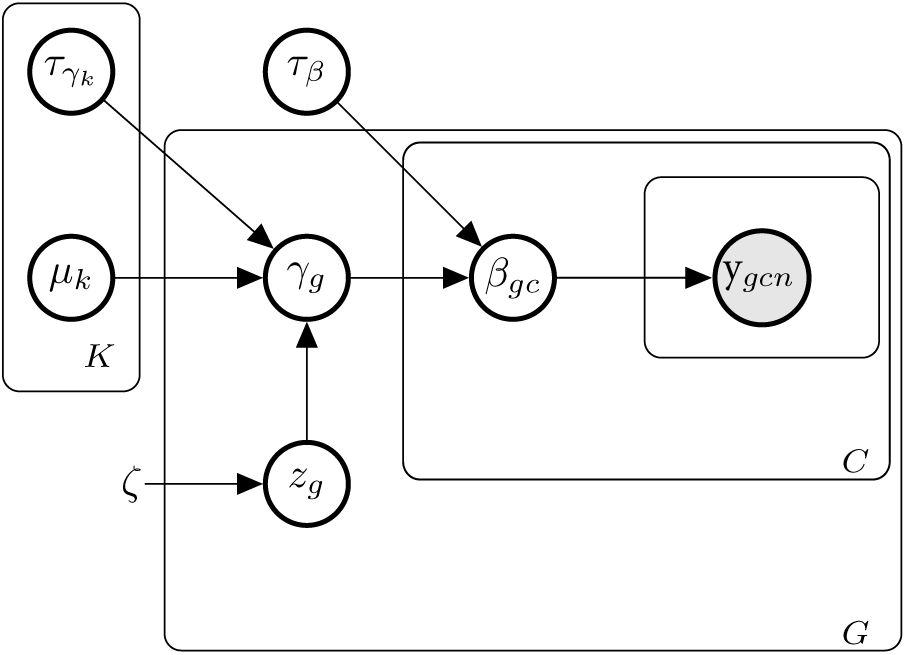
The basic structure of an SHM is a combination of a HM and a MRF. On the first level of the hierarchy of the HM we model the data *y*_*gcn*_ for a gene *g*, condition *c* and observation *n* as an exponential family distribution that is parameterized by a gene-condition effect *β*_*gc*_. The gene-condition effect is modelled as a random variable with parameters *γ*_*g*_, the gene effect, and a nuisance parameter *τ*_*β*_. The gene effects are parameterized by a MRF *z*_*g*_ which clusters the latent gene effects, and vectors of means 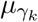 and standard deviations 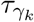.

The top level of the hierarchy is now the categorical MRF from Equation (2.2) (abbreviated for notational simplicity). We replace the uni-variate distributions of *µ* and 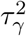 in Equation (2.1) with *K*-dimensional distributions. The MRF induces a clustering in the latent space of *γ* and not in the data space **y**. We are primarily interested in grouping genes, i.e., the variables *γ* and not the actual data, i.e., the effect sizes of the replicates of single interventions *y*_*gcn*_. In addition, clustering in the data space is not only too difficult due to the noisy readouts of biological interventions, but also hardly ever of biological interest. The rest of the model stays as in (2.1).

The SHM compensates misspecification or incomplete domain knowledge by pushing down biological prior knowledge using the top-level MRF through the different hierarchies, thereby informing inference of latent parameters *γ*_*g*_ and *β*_*gc*_. More specifically, either the real data generating process of the biological system under study is unknown and not all covariables and confounders that take influence are measured or known, or we cannot include them as covariables, as inclusion would lead to ill-defined models that are not, or only weakly, identifiable, e.g., in low sample size settings.

The model in Equation (2.4) serves as a basic structure and needs to be adjusted for specific domains, i.e., complemented with other covariables and appropriate distributions for the data at hand.

## 3. Models for genetic perturbation data

We use the backbone of the SHM (2.4) to build a concrete model for data from genetic perturbation screens. This requires specifying appropriate prior distributions for variance components and a distribution for the data, as well as supplementing the data generating process with appropriate covariables. For multi-condition genetic perturbation data sets we define the model as follows:

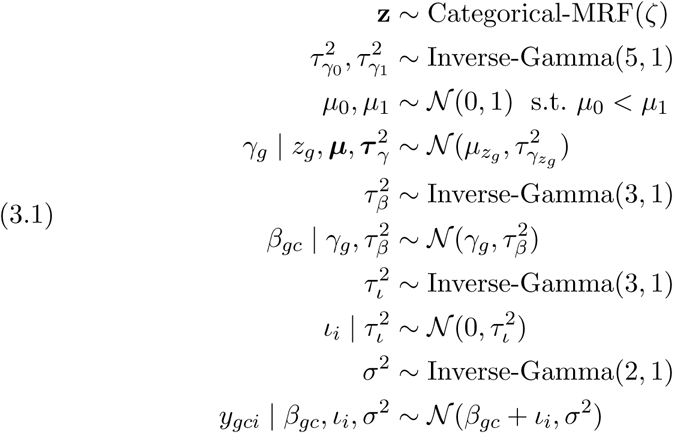

where we included variable *ι*_*i*_ to correct for effects an intervention might have, e.g., cytotoxicity of the gRNA used for perturbation (note that this introduces another level of hierarchy for the interventions *i*). We choose inverse-Gamma distributions as priors for variance components, following Kass et al. (2006). The rest of the model is the same as in Equation (2.4).

Following (Meyers et al., 2017), if information of copy number aberrations *c*_*gc*_ is available, we include it as covariate for every gene and condition. Adjusting for copy number aberrations is necessary, because in regions of high copy number gain interventions can occur multiple times leading to a DNA damage response and cell cycle arrest. This in turn leads to no proliferation and observed readouts are low independent of the gene that has been perturbed. This part of the model then becomes

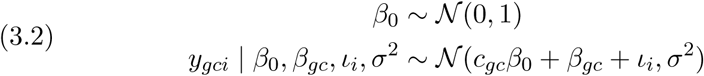

where the parameters that are not endowed with priors are distributed as in model (3.1).

## 4. Model criticism

We evaluate our model using several simulated data sets and a biological data set from the DepMap portal to show that the inference of parameters can indeed be improved using biological prior knowledge. We compare SHMs to models, where we replace the MRF on top of the hierarchy of Equation (2.4) with a conventional mixture model leaving the rest of the model identical. In this way we can assess the influence of the network itself.

We investigate the SHMs described in Equations (3.1) and (3.2), and are interested in estimating effect sizes *γ*_*g*_ and classifying genes as essential (*z*_*g*_ = 1) or non-essential (*z*_*g*_ = 0), since these are the parameters of greatest biological importance. Thus we use *K* = 2 mixture components. Every gene *g* is perturbed using multiple different gRNAs *i* in multiple conditions *c*. If a gene is essential, i.e, *z*_*g*_ = 1, and has a negative gene effect, *γ*_*g*_ *<* 0, we expect to observe cell death upon intervention, while we expect no change in cell proliferation when the gene is non-essential and has an effect size *γ*_*g*_ ≈ 0.

We first introduce criteria to evaluate the models, and then apply different SHMs to the data sets.

### 4.1. Criteria of model criticism

To evaluate the model, we are primarily interested in the estimates of the latent variables ***γ*** and **z**. We assess the accuracy of the inference of the posterior of gene effects by comparing their estimated posterior means 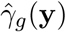 to their true values *γ*_*g*_ using squared error loss

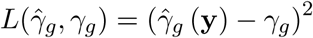

In order to assess the correct inference of posterior labels, we compare means of posterior labels 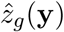 to their true values *z*_*g*_ using the number of correct and incorrect classifications. Specifically, we estimate the number of true positives, i.e., the number of predictions of correctly classifying a gene as

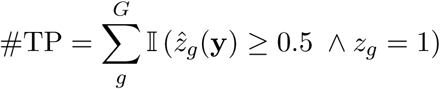

and the number of false negatives as

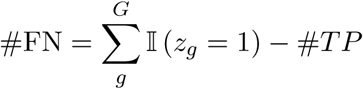

We evaluate the number of false positive inferences as

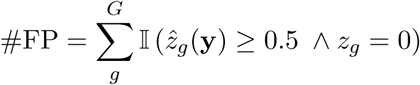

and the number of true negatives analogously as

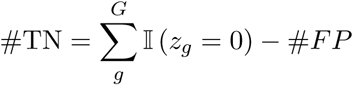

We follow the hypothetico-deductive model of (Gelman and Shalizi, 2013; Gelman et al., 2013) to evaluate posterior inference visually by computing posterior predictive distributions (PPCs) and plotting them as density estimates over the data Gabry et al. (2019). The predictive posterior is computed as

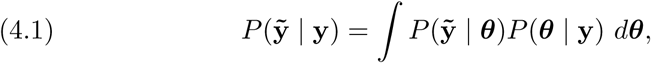

where ***θ*** = (**z**, ***γ***, …) is a vector of all random variables of an SHM.

### 4.2. Simulated data

We first evaluate our model on multiple simulated data sets in order to show that including graph prior knowledge can indeed inform posterior inference when the data generation process is misspecified. Specifically, we compare the SHM in (3.1) to a model that uses the same hierarchical structure, but replaces the MRF (2.2) with a conventional mixture model, i.e., we only replace the first line in (3.1) with:

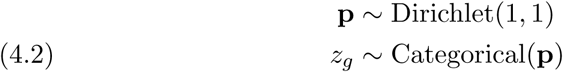

Since the HM part is the same in both models, any differences in the inference of posteriors can only be explained through the MRF, or mixture model, respectively. We briefly describe the different steps how we generated the data sets for the validations:

1. Generate a scale-free network of genes using the `powerlaw_cluster_graph` function from `networkx` package (Hagberg, Schult and Swart, 2008).
2. Separate the genes of the networks into two groups, a group of essential genes and a group of non-essential genes.
3. For essential genes sample *γ*_*g*_ ∼ 𝒩 (−1, 0.25), for non-essential genes sample *γ*_*g*_ ∼ 𝒩 (0, 0.1) (argueing that genes with no effect should have a lower variance, since no effect should be measured).
4. Sample *β*_*gc*_ ∼ 𝒩 (0, 0.25).
5. Sample *ι*_*i*_ ∼ 𝒩 (0, 0.1).
6. In order to simulate low-quality interventions *i*, set gRNA activity of some of the essential genes to *a*_*i*_ = 0.1 (we call these genes in the following essential genes with low affinity gRNA). For the other gRNAs we assume high-quality interventions setting *a*_*i*_ = 1.
7. Sample *y*_*gci*_ ∼ 𝒩 (*a*_*i*_*β*_*gc*_ + *ι*_*i*_, *σ*^2^) with *n* = 10 replicates for every combination of *g, c* and *i* and noise variances *σ*^2^ ∈ {0.1, 0.2, 0.3, 1}.

Hence, in order to simulate misspecification, i.e., in this case incomplete domain knowledge, we use a data generating process that includes a covariate for low-quality interventions *a*_*i*_, but then model the simulated data without it. Incomplete domain knowledge is frequent in biology.

For the first validation, we were interested in the effect of low-activity gRNAs of essential genes when neighborhoods consist entirely of other essential genes (Figure 3a). For perfect interventions, i.e., when all affinities *a*_*i*_ = 1 and the data generating processes are perfectly captured by models (3.1) and (4.2), the two models are identical, as no information of the labels of neighbors is needed, and hence inference should be the same (Figure 3b top). The posterior distributions of the gene effects *γ*_*g*_ and class assignments *z*_*g*_ of these genes illustrate this (Figure 4 top). However, in the case of some gRNAs having low affinities the influence of the neighbors, mediated through the MRF, improves inference substantially (Figure 3b bottom; Figure 4 bottom).

**Fig 3:**
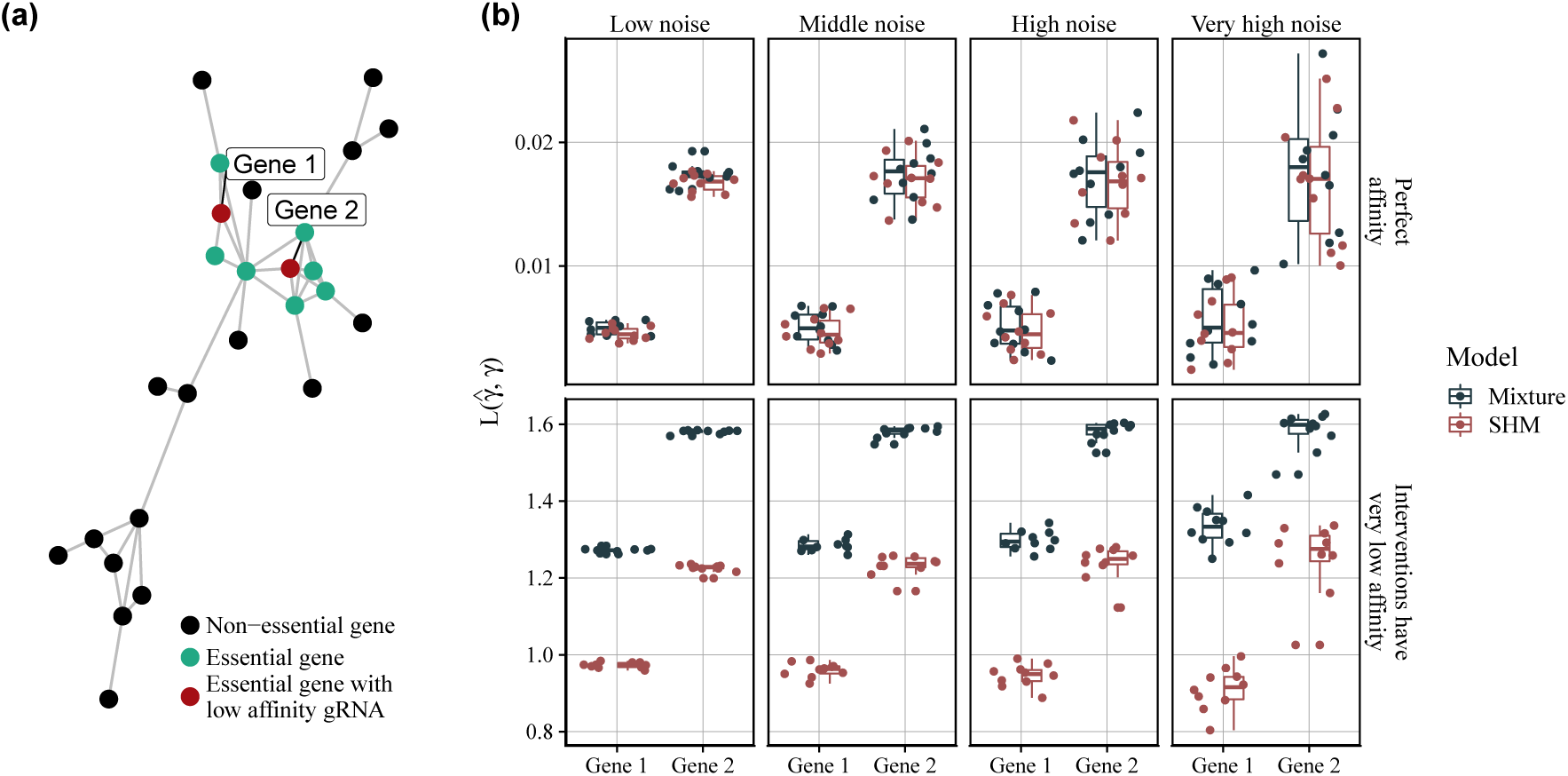
Benchmarking of parameter inference for genes that are embedded in neighborhoods of genes with the same label. (a) The network shows groups of essential genes (green/red) and non-essential genes (black). For red essential genes we simulated data where some gRNAs have low activity, i.e., where interventions do not work as intended and where the data does not show that the genes are essential. (b) Performance of the mixture model and the SHM. Every box shows the results of 10 simulated data sets and different noise levels. The y-axis shows the squared error loss of posterior means of gene effects of gene 1 and 2 (lower is better). If data are generated with perfect activity for the red nodes, the mixture and the SHM perform equally good as expected (top row). If we generate gRNAs with low activity for the two red nodes the SHM clearly outperforms the conventional mixture model (bottom row).

**Fig 4:**
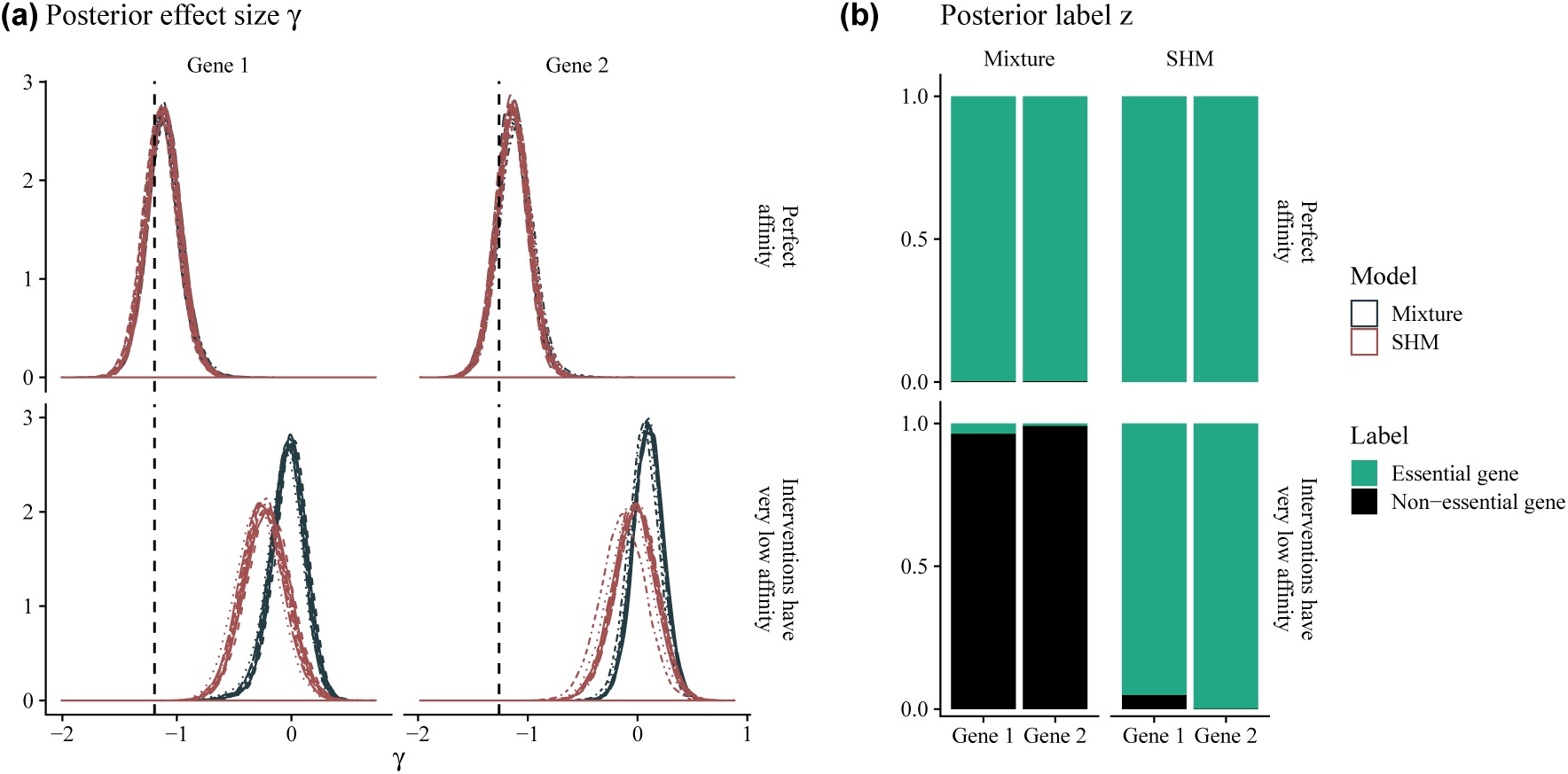
Posterior distributions of the effect sizes *γ*_*g*_ and labels *z*_*g*_ for genes that are embedded in neighborhoods of genes with the same label. The experimental setting is as in Figure 3, i.e., the groups of the graph are the same and the red nodes have low-activity gRNAs. (a) As expected the SHM and the mixture model infer the same posterior distributions when all interventions work as intended (top row). When some of the gRNAs have low activity, the SHM moves the posterior distribution closer to the true value (dashed line) and outperforms the mixture model (bottom row). (b) Both models infer the posterior probabilities of the labels *z*_*g*_ correctly when gRNAs work as intended (top row). In the case of low-affinity gRNAs, the SHM is still able to infer the correct labels, while the mixture model fails to do so (bottom row).

We then assessed the influence of the MRF on inference when essential genes with low-affinity gRNAs have essential as well as non-essential neighbors (Figure 5a). In this case, as long as the essential genes with low affinities have more essential neighbors than non-essential ones, estimation of parameters improves. If there are more non-essential neighbors than essential ones, the mixture model has negligibly better performance than the SHM. Vice versa, if there are more essential neighbors, the SHM outperforms the mixture model. In general assessing settings where the number of essential and non-essential neighbors is roughly the same is difficult, because posterior inference will be dominated by the data itself and not the MRF, i.e., the effect of the MRF vanishes, because the influence of essential and non-essential neighbors is equally strong and cancels out (Figure 5b). In these settings, both models achieve similar results.

**Fig 5:**
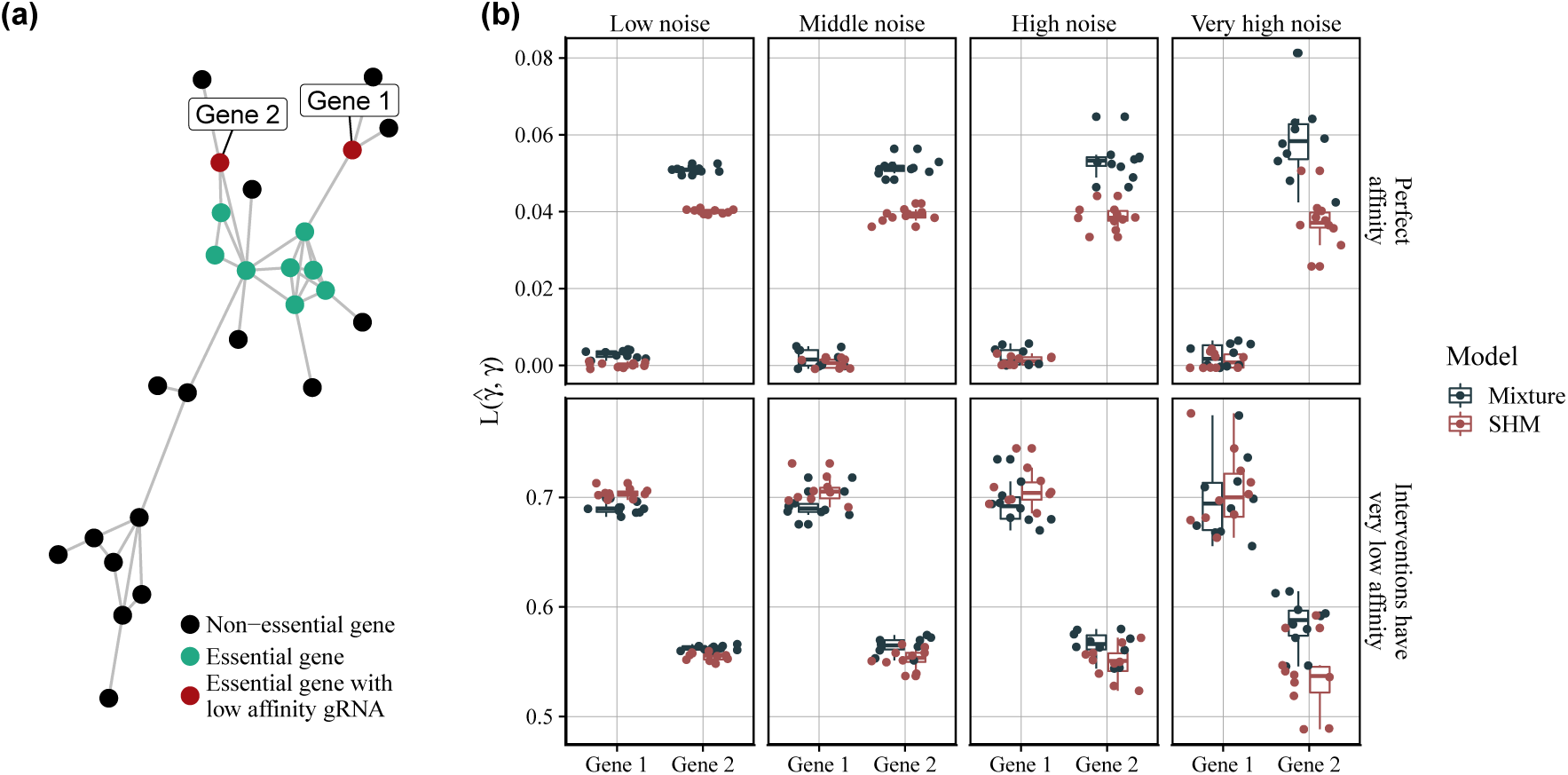
Benchmarking of parameter inference when the number of neighbors with the same label is one higher, or lower, than non-essential neighbors. (a) The network shows the same graph of genes as in Figure 3 with the difference that now red genes belong to a different neighborhood. (b) When the number of essential and non-essential neighbors is roughly the same, the influence of the SHM vanishes and the data primarily informs inference. Hence, it is difficult to assume that the SHM outperforms the mixture model, even though the performances are improved for gene 2. For gene 1 the mixture model slightly outperforms the SHM. Generally, in scenarios where the number of essential and non-essential neighbors is the same, both models should perform similarly.

Finally, we show how one can break our model, namely by embedding an essential gene with low-affinity gRNAs in a neighborhood of exclusively non-essential genes (Figure 6). The neighborhood then wrongly informs the inference of the gene’s class assignment and the mixture model outperforms the SHM. In practice, the situation is unlikely to occur, because genes that are neither functionally nor biochemically related do not form meaningful protein-protein interactions and are consequently not connected with edges but instead found in different sub-modules of the graph.

**Fig 6:**
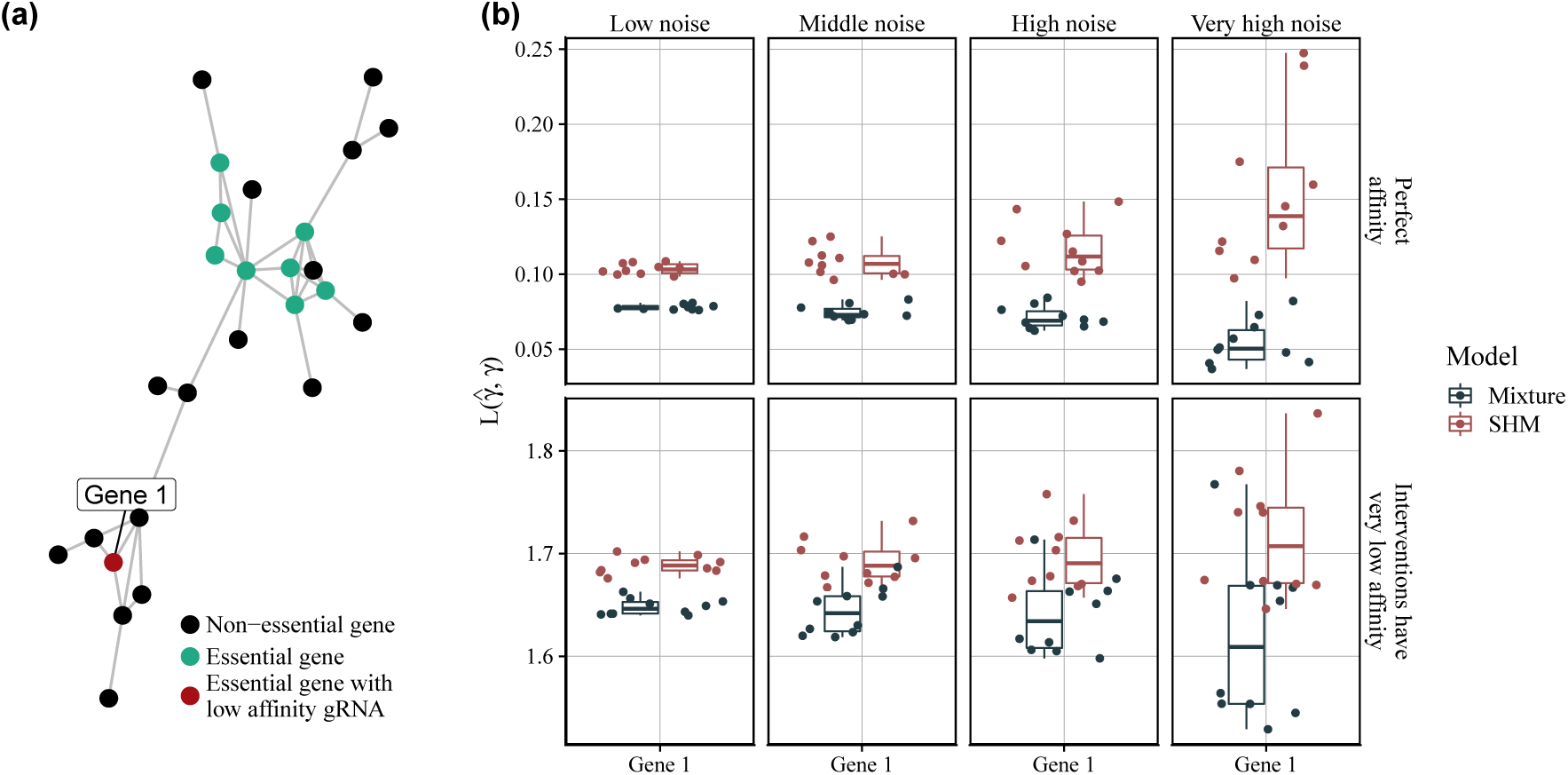
Benchmarking of parameter inference for genes that are embedded in neighborhoods of genes with a different label. (a) The network shows the same graph of genes as in Figure 3 with the difference that now red essential genes are surrounded by non-essential genes. (b) In this scenario the SHM wrongly informs posterior inference and the estimates of the effect sizes are worse in comparison to the mixture model, both for gRNAs with perfect affinity, as well as for gRNAs with lower affinity.

### 4.3. Biological data

We applied the SHM to a biological data set form the DepMap portal (Tsherniak et al. (2017); Meyers et al. (2017); data downloadable from Broad Institute (2019)). The DepMap data consist of several cancer cell lines (conditions) for which genetic perturbation experiments using CRISPR have been conducted and the effects of a knockout on cell proliferation have been measured. If a gene is essential we expect to observe cell death upon intervention, while we expect no change in cell proliferation when the gene is not essential.

We downloaded the data from the the DepMap portal and selected screens with *n* = 4 replicates and a sufficiently high Cas9 activity, leaving a total of 7 experiments (Table 1).

**TABLE 1.** Meta data of the biological data set from the DepMap portal.

| ID | Cell-line | Disease | Disease subtype | Set |
| --- | --- | --- | --- | --- |
| ACH-000120 | CHP212 | peripheral nervous system | neuroblastoma | train |
| ACH-000414 | NCIH1944 | lung | lung NSC | train |
| ACH-000841 | NCIH2087 | lung | lung NSC | train |
| ACH-000312 | SKNBE2 | peripheral nervous system | neuroblastoma | test |
| ACH-000226 | SUPM2 | lymphoma | ALCL | test |
| ACH-000311 | NCIH2122 | lung | lung NSC | test |
| ACH-000118 | HUPT3 | pancreas |  | test |

The DepMap data provides information about true essential and non-essential genes from (Hart et al., 2014, 2015), i.e., information on some of the genes regarding their label. In order to be able to validate our estimates, i.e. whether we classified genes correctly, we reduced the set of genes to these controls, leaving roughly 500 genes. We used the SHM (3.2) to model the data.

The latent labels **z** are distributed as in Equation (2.2) with edge potentials as defined in Equation (2.3) and constant weights *w*_*ij*_ = 1. We downloaded the String PPI network (Szklarczyk et al., 2018) and removed all edges with low confidence, i.e., edges with a score of less than 500 (Figure 7). This step is done in an effort to reduce potential false positive edges, i.e., edges erroneously inferred using yeast two-hybrid (Y2H) or other technologies to detect PPIs. In the network, essential genes and non-essential genes seem to be surprisingly well separated. Curious about this finding, we compared this network with the functional network from (Wu, Feng and Stein, 2010) and the physical interaction network from (Oughtred et al., 2018) and found that both groups of genes were equally well separated in these networks, although not all genes from the DepMap data were contained in them, which possibly explains the high number of singletons in these networks (Supplementary figure 1).

**Fig 7:**
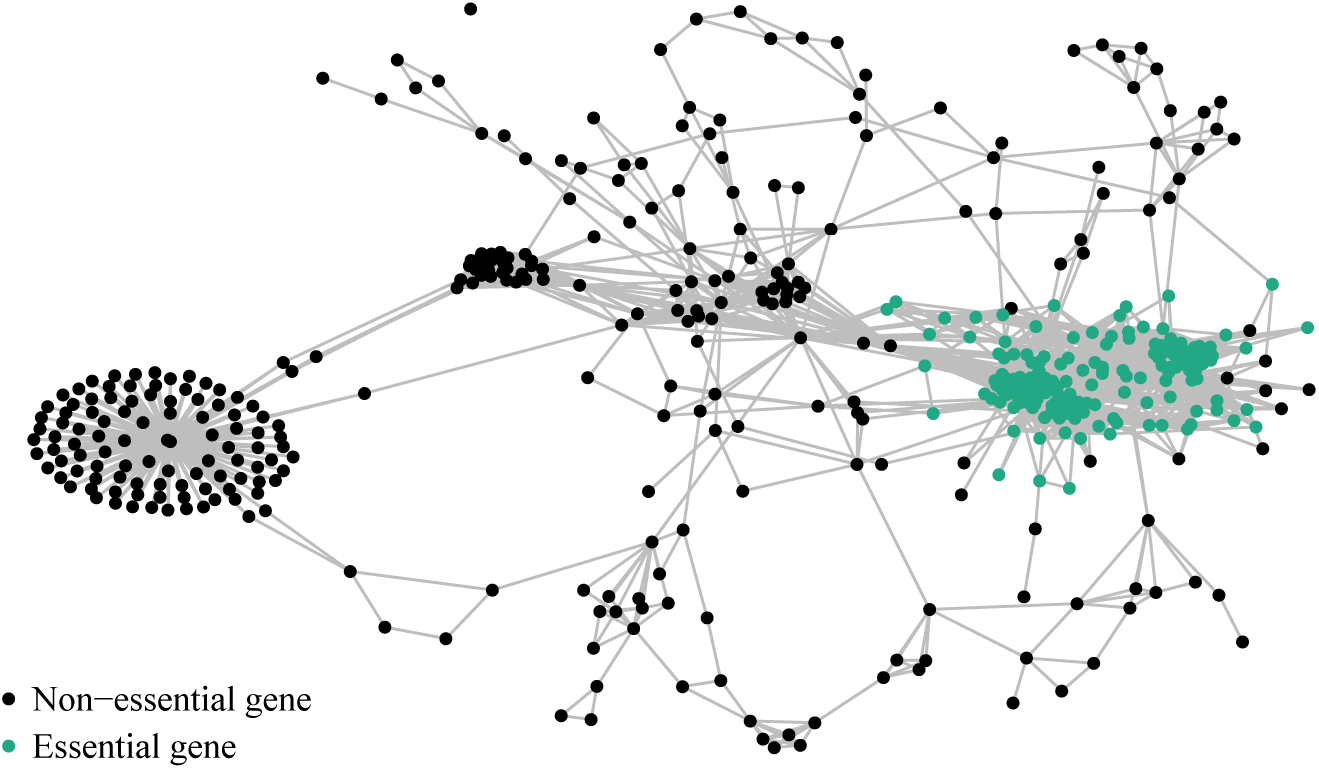
Biological network used for inference of essential genes from the DepMap data. In order to apply the SHM to real data, we downloaded the STRING network that encodes gene-gene interactions. For visualization we also downloaded information about gene essentiality and colored the nodes respectively. Essential genes seem to cluster into modules of high-connectivity (compare Supplementary figure 1).

Since the DepMap portal provides estimates of the activity of gRNAs (Doench-Root scores), i.e., quantification how well a gRNA works, we fit a second model that has an additional covariate for gRNA affinity *a*_*i*_, but is otherwise identical to (3.2). For most other experiments, estimates of gRNA affinity are not available and we were only able to include it here. The data generating process then changes to

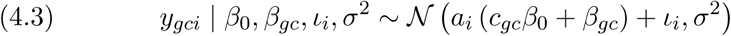

We first estimated the hyperparameter *ζ* using a grid search in [0.05, 0.1, …, 1] with training data consisting of three cell lines (train data sets in Table 1) resulting in an optimal estimate of *ζ* = 0.15. We then inferred the parameters of the models using test data of four cell lines (Table 1). As in the simulated data benchmarks, we compared the model against a model where the MRF is replaced with a conventional mixture model (Equation 4.2) with two components. As before, with this approach differences in the parameter estimates are due to the MRF alone, and not the HM. We are assessing in total four different models (Table 2).

**TABLE 2.**
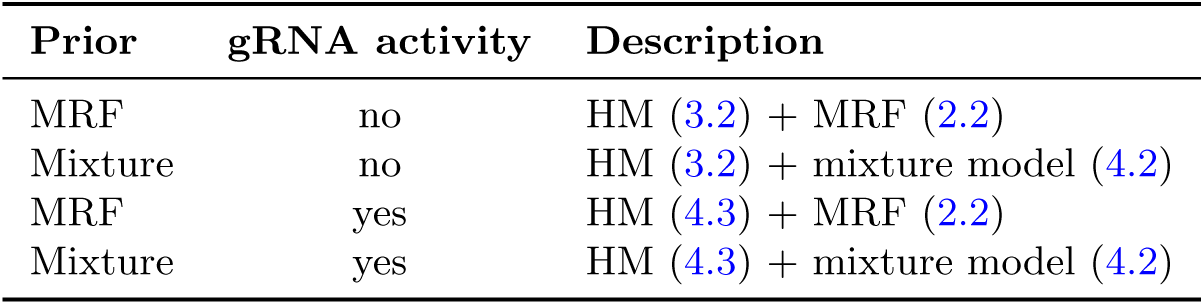
Overview of different models used for model criticism with biological data.

| Prior | gRNA activity | Description |
| --- | --- | --- |
| MRF | no | HM (3.2) + MRF (2.2) |
| Mixture | no | HM (3.2) + mixture model (4.2) |
| MRF | yes | HM (4.3) + MRF (2.2) |
| Mixture | yes | HM (4.3) + mixture model (4.2) |

We found that in both cases biological prior information improved inference of the posterior labels (Figure 8). While the models with Doench-root scores are very similar and the improvement of the SHM is only marginal, reducing the number of false negatives from 8 to 2, the improvement of the SHM over the clustering model when not including the Doench-root score is substantial: we reduce the number of FNs from 41 to 1. Interestingly we do not observe improvement in false positives (all 2), which emphasizes our initial hypothesis: inference of essential genes is mainly a problem of high false-negative rates and not of false-positive rates. Between the SHM models there are hardly any improvements, underlining our hypothesis that better, more robust inferences form noisy data sets can be made when network information is incorporated. On the other hand, the results of the mixture models are very different which emphasizes the fact that careful model building is required for interventional studies, a problem that gets aggravated with missing domain knowledge.

**Fig 8:**
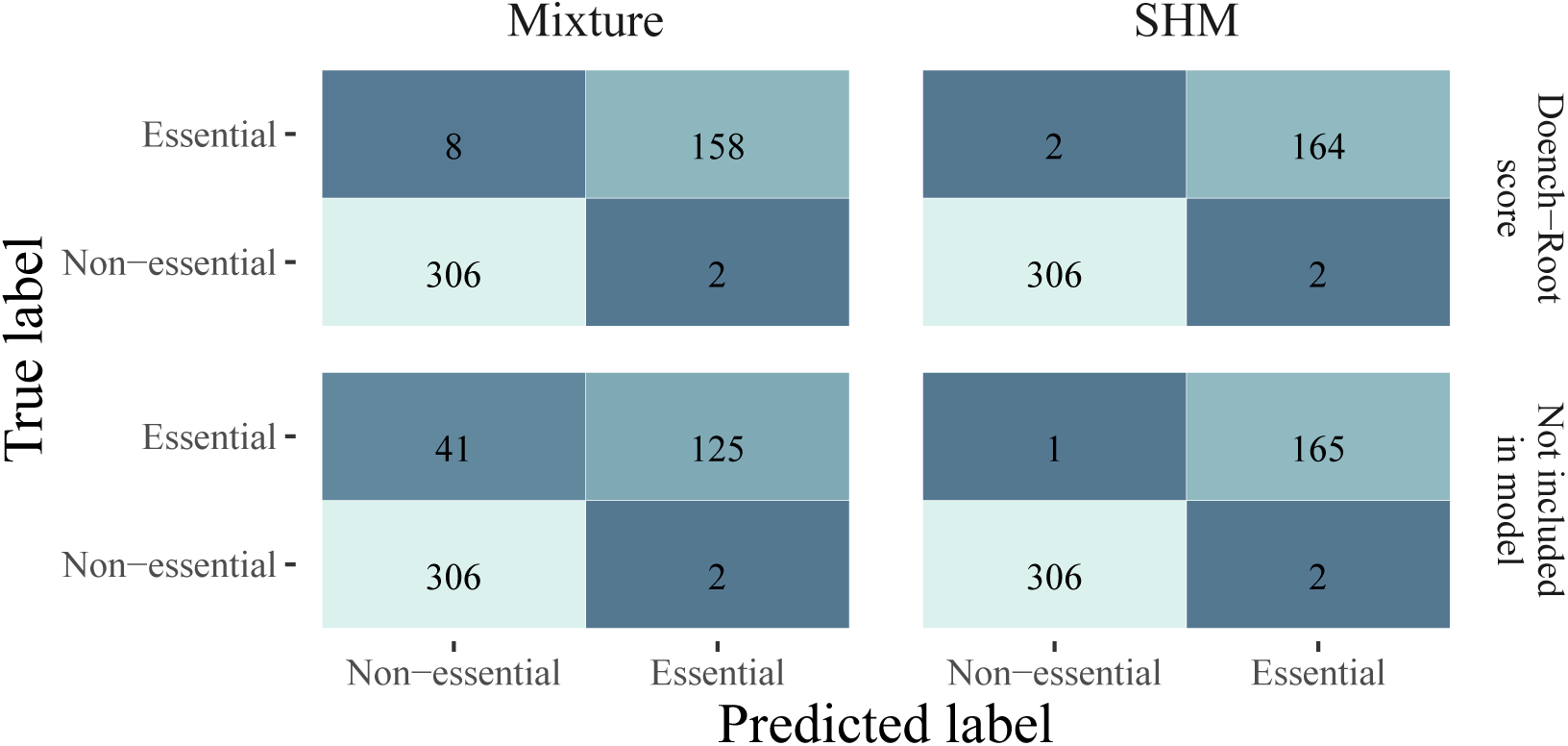
Confusion matrices of true and estimated essential genes of four biological models inferred from the DepMap data. Every confusion matrix shows the number of false positives and false negatives for each model (Table 2). The SHM outperforms the mixture models if Doench-Root scores, i.e., covariables quantifying gRNA activity, are included by reducing the number of false negatives from 8 to 2 (top row). When Doench-Root scores are not included, the number of false negatives is reduced from 41 in the mixture to 1 in the SHM (bottom row).

Considering the posterior predictive distributions of the four models that we fit, we found the distributions to be almost identical (Figure 9). This shows that the main differences are mainly in inferring the nested parameters, while the actual data generating process stays the same.

**Fig 9:**
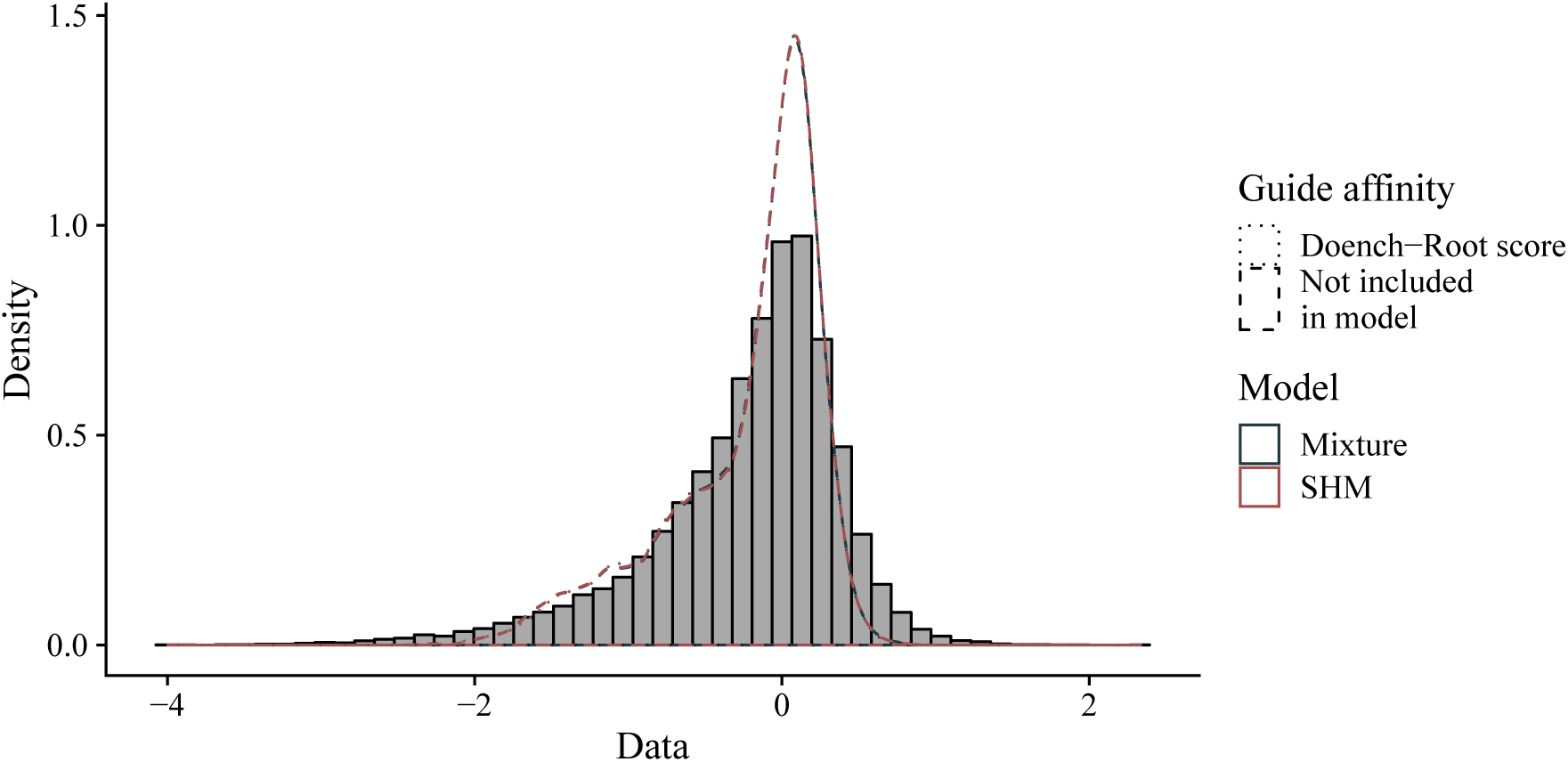
Posterior predictive distributions (PPCs) of the DepMap genetic screening data sampled using four biological models (Table 2). We sampled data from the posterior predictive distributions using Equation 4.1 and overlayed the histogram of the original biological data set with the kernel denisty estimates of the sampled data. All four models can adequately predict new data from the posterior distributions. The density curves are close to the original data, especially in the lower tail of the histogram. The PPCs underestimates the upper tail of the histogram slightly.

## 5. Implementation

The graphical model representation of the SHM in (2) has a tree structure (Figure 2) which allows for Gibbs sampling to infer posterior distributions. In particular, we developed a custom Metropolis-within-Gibbs sampler using the probabilistic programming language `PyMC3` (Salvatier, Wiecki and Fonnesbeck, 2016). Here, we show how one can simulate from the posterior distribution of the SHM in Equation (2.4). The sampler can be easily extended to more complex models with more covariates and latent variables, as long as the Markov blanket of the class labels **z** stays the same.

We denote by *D* the data and 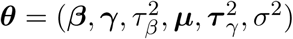 the continuous parameters of the model. The full posterior of Equation (2.4) has the following form:

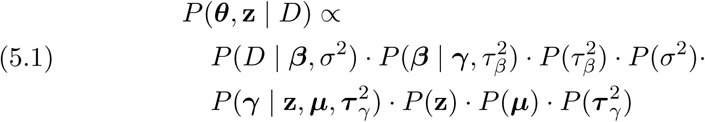

For every iteration *t*, a sample (***θ*, z**)^*t*^ from the full posterior using Metropolis-within-Gibbs can be obtained in two steps:

1. draw **z**^*t*^ from *P* (**z** | *D*, ***θ***^*t*−1^) ∝ *P* (*γ* | **z, *µ***, *τ*_*g*_) · *P* (**z**) using Gibbs sampling,
2. draw ***θ***^*t*^ from *P* (***θ*** | *D*, **z**^*t*^) using Hamiltonian Monte Carlo.

Developing the Metropolis-within-Gibbs sampler with `PyMC3` only requires implementing a custom sampler for the Markov random field **z** in step one (Appendix A for more details). For the continuous variables ***θ*** we use a Hamiltonian Monte Carlo (HMC) (Neal, 2011; Betancourt, 2017) variant, the No-U-Turn sampler (Hoffman and Gelman, 2014), provided by `PyMC3`. In order to avoid divergences and for efficiency, we use a non-centered parameterization whereever possible (Betancourt and Girolami, 2015).

We implemented the sampler described above in a Python package which is available from GitHub at github.com/cbg-ethz/shm. The GitHub repository also contains the implementations for the models used in section 4.

## 6. Discussion

We introduced a new family of models, the *structured hierarchical model* (SHM), which combines Bayesian hierarchical models with categorical Markov random fields. Through the random field, it is possible to aid posterior inference by probabilistically incorporating biological network information into the model. Since the random field introduces a latent categorical variable **z** which is used to parameterize the hierarchical model, we induce a mixture model of the top-level latent variables of the hierarchical model. In our application of perturbation data, the clustering allowed labelling genes as essential or non-essential.

We applied the SHM to simulated data as well as a biological data set from the DepMap portal and found that including biological prior knowledge generelly improves estimation of parameters. We found that the biological networks we considered were all very similar in grouping functionally related genes together in a biological meaningful way.

The SHM is especially useful for noisy data sets consisting of multiple conditions which are common in interventional, biological studies. We expect our model to be of use for computational biologists, because it demonstrates how to incorporate biological networks into probabilistic analyses. For the inference of parameters, HMs borrow statistical strength from other hierarchy levels. Additionally, when choosing appropriate priors, Bayesian inference has an auto-regularizing effect which makes it applicable even for data sets with low sample size. Consequently, Bayesian HMs are very useful in different branches of computational biology, powerful tools due to their statistical properties, and easily interpretable.

However, the SHM also has some drawbacks. In general, full Bayesian inference using sampling is computationally highly demanding and can be difficult, for instance on a genome-wide scale. Sampling is further complicated due to the high-dimensional categorical variable **z**, because sampling discrete parameters usually yields low effective sample sizes and thus requires long chains for convergence. While we usually marginalize out **z** in conventional mixture models, this cannot be done here due to the conditional dependencies encoded in the MRF. Furthermore, not much research has focussed on the theoretical and practical properties of combining an HMC for continuous parameters with a Gibbs sampler for discrete ones. In order to speed up posterior inference, approaches using variational inference could be adopted, but would require to either continuously relax or reparameterize the discrete Markov random field **z**, e.g., as in (Jang, Gu and Poole, 2017) or (Guo and Schuurmans, 2006). Applying reparameterization to conditionally dependent random variables has not received much attention by researchers though. Thus, a natural, non-Bayesian approach could be *maximum a posteriori* (MAP) inference, or approximate approaches such as empirical Bayes (Efron, 2012). MAP inference, however, has the obvious drawback of not yielding uncertainty estimates.

The SHM described in Equation (2.4) represents a basic structure which needs to be adjusted to specific domains. Hence, we did not specify priors for variance parameters. In our analyses above, the use of inverse-Gamma distributions for variance parameters has no real justification other than being weakly-informative and conditionally conjugate. While for the sake of demonstration of the SHM this is acceptable, in general, one can follow principled ways for prior specification, such as (Gelman, 2006; Gelman, Simpson and Betancourt, 2017; Kass and Wasserman, 1996).

We hope our contributions will be only a first step towards integrating biological prior knowledge into probabilistic models and that more research is invested into the topic in general and into efficient (variational) approximations in particular.

## APPENDIX A: GIBBS SAMPLING A CATEGORICAL MRF

Denote *D* the data and ***θ*** = (***β***, *τ*_*β*_, *σ*^2^) (note that we use different notation for ***θ*** as before). The first step of the Metropolis-within-Gibbs sampler draws the posterior 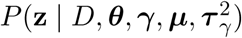. Following Wainwright and Jordan (2008), a sample is obtained by conditioning on the Markov blanket of **z**, which is:

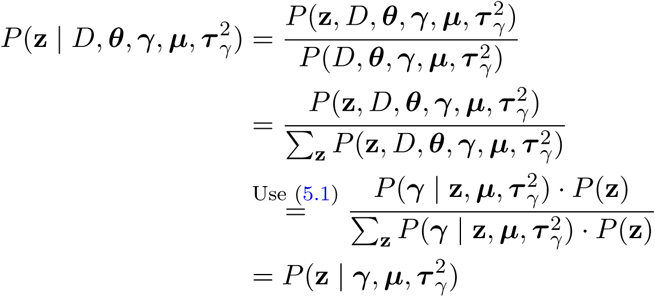

We sample from 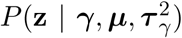 componentwise for every *z*_*i*_ ∈ {1, …, *K*}, *i* ∈ {1, …, *G*} using a Gibbs-sampler:

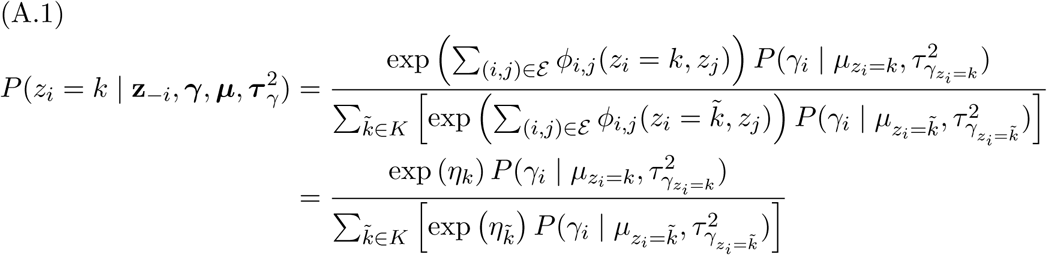

where we introduced *η*_*k*_ = Σ(*i,j*)_∈*ε*_ *ϕ*_*i,j*_ (*z*_*i*_ = *k, z*_*j*_) for brevity. In the binary case, Equation A.1 can be simplified to

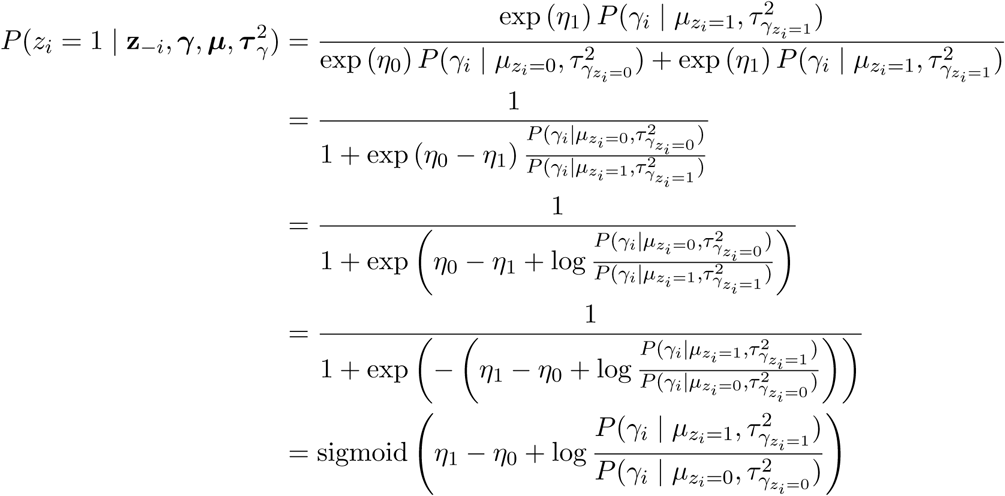

## SUPPLEMENTARY MATERIAL

### Supplementary material: Figures

(link and doi to be assigned by journal). The supplementary material contains figures of other biological networks and the linear dependency of readouts and copy number amplifications.

