## Supplementary material for "Structured hierarchical models for probabilistic inference from perturbation screening data"

**Supplementary figure 1. Other biological networks.** Shown are the functional network from (Wu et al., 2010) and the BioGRID network (Oughtred et al., 2018). Both networks have similar modules as the network from STRING (Szklarczyk et al., 2018), where essential genes are clustered in high-density regions. However, both networks are missing several genes from the data from the DepMap portal (Tsherniak et al., 2017; Meyers et al., 2017; Broad Institute, 2019), which probably explains the high number of singletons in comparison to STRING.

(a)

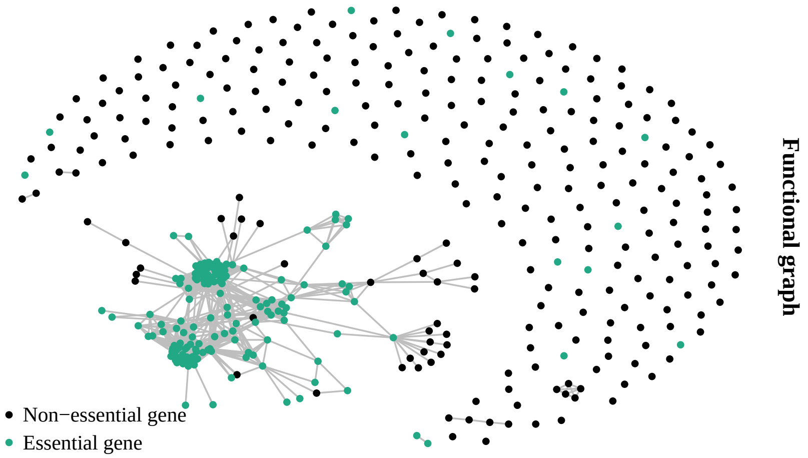


(b)

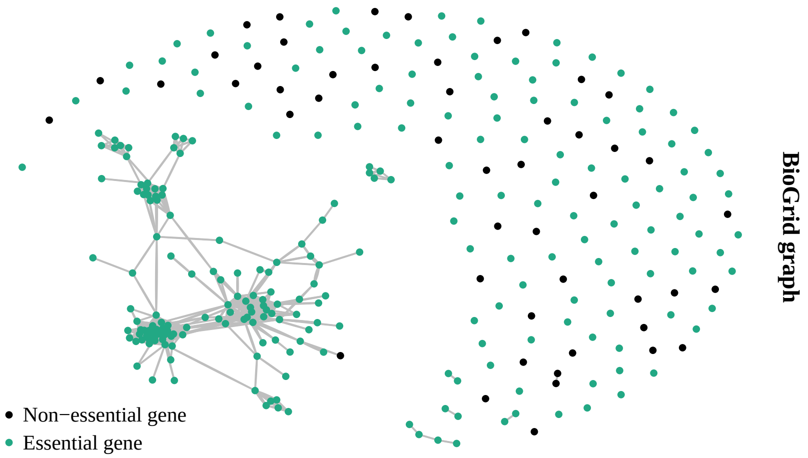

**Supplementary figure 2. Copy numbers amplifications.** The plot shows the copy number alterations against the observed readouts. For the cancer data that we used (Tsherniak et al., 2017; Meyers et al., 2017; Broad Institute, 2019), we could not observe the same impact of copy number alterations on the readout as described in Meyers et al. (2017); Aguirre et al. (2016).

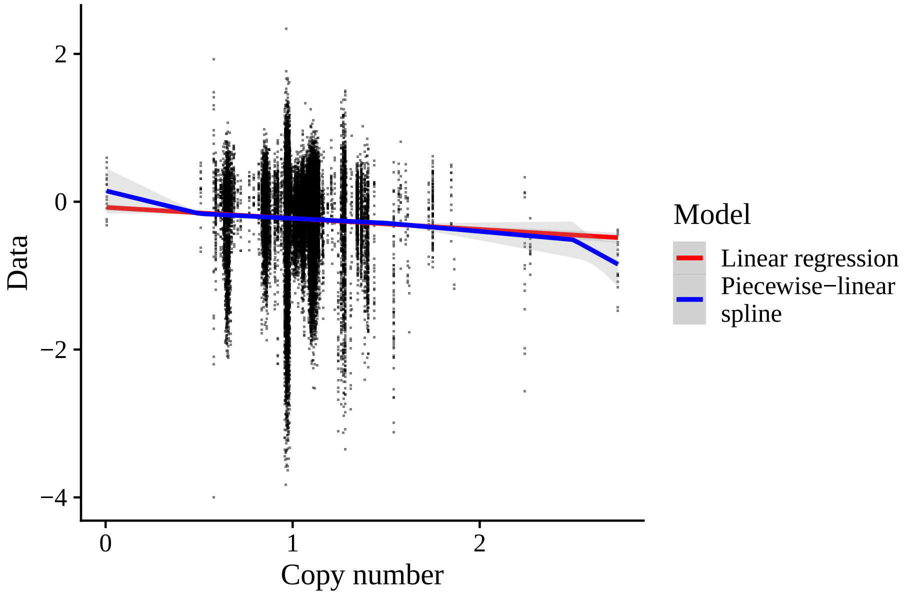


### References.


Andrew J Aguirre, Robin M Meyers, Barbara A Weir, Francisca Vazquez, Cheng-Zhong Zhang, Uri Ben-David, April Cook, Gavin Ha, William F Harrington, Mihir B Doshi, et al. Genomic Copy Number Dictates a Gene-Independent Cell Response to CRISPR/Cas9 Targeting. *Cancer Discovery*, 6(8):914–929, 2016.

Broad Institute. DepMap Achilles 19Q1 Public. Fileset on figshare, 2019. URL `doi:10.6084/m9.figshare.7655150`.

Robin M Meyers, Jordan G Bryan, James M McFarland, Barbara A Weir, Ann E Sizemore, Han Xu, Neekesh V Dharia, Phillip G Montgomery, Glenn S Cowley, Sasha Pantel, et al. Computational correction of copy number effect improves specificity of CRISPR–Cas9 essentiality screens in cancer cells. *Nature Genetics*, 49(12):1779–1784, 2017.

Rose Oughtred, Chris Stark, Bobby-Joe Breitkreutz, Jennifer Rust, Lorrie Boucher, Christie Chang, Nadine Kolas, Lara ODonnell, Genie Leung, Rochelle McAdam, et al. The BioGRID interaction database: 2019 update. *Nucleic acids research*, 47(D1):D529–D541, 2018.

Damian Szklarczyk, Annika L Gable, David Lyon, Alexander Junge, Stefan Wyder, Jaime Huerta-Cepas, Milan Simonovic, Nadezhda T Doncheva, John H Morris, Peer Bork, et al. STRING v11: protein–protein association networks with increased coverage, supporting functional discovery in genome-wide experimental datasets. *Nucleic Rcids Research*, 47(D1):D607–D613, 2018.

Aviad Tsherniak, Francisca Vazquez, Phil G Montgomery, Barbara A Weir, Gregory Kryukov, Glenn S Cowley, Stanley Gill, William F Harrington, Sasha Pantel, John M Krill-Burger, et al. Defining a Cancer Dependency Map. *Cell*, 170(3):564–576, 2017.

Guanming Wu, Xin Feng, and Lincoln Stein. A human functional protein interaction network and its application to cancer data analysis. *Genome Biology*, 11(5):R53, 2010.
